# Commonly prescribed medicines antagonise anti-MRSA antibiotics and select for resistance

**DOI:** 10.64898/2026.03.31.715408

**Authors:** Zoha Sohail, Henry A. Claireaux, Andrew M. Edwards, Edward J.A. Douglas

**Affiliations:** Centre for Bacterial Resistance Biology, Imperial College London, London SW7 2AY, United Kingdom; Department of Infectious Disease, Imperial College London, London SW7 2AY, United Kingdom; Department of Infectious Disease, King’s College London, London, United Kingdom; Academic Department of Military Trauma & Orthopaedics, Research & Clinical Innovation, ICT Centre, Birmingham, United Kingdom

## Abstract

Many commonly prescribed non-antibiotic medicines have off-target antimicrobial activity, yet their impact on antibiotic efficacy remains poorly understood. In this study, we investigated eight widely used UK prescription medicines and identified simvastatin, amlodipine, and fluoxetine as growth inhibitory towards methicillin-resistant *Staphylococcus aureus* (MRSA). These drugs disrupt bacterial membranes, with amlodipine and fluoxetine also triggering stress responses linked to cell wall and membrane damage. Further mechanistic analysis using transposon mutant screening revealed that simvastatin impairs cell wall synthesis by inhibiting the mevalonate pathway. Notably, checkerboard assays demonstrated antagonistic interactions: simvastatin reduced the efficacy of β-lactams and vancomycin, amlodipine with vancomycin and daptomycin, and fluoxetine with vancomycin activity. Prolonged exposure to these drugs also accelerated resistance development to vancomycin and daptomycin. Together, these findings underscore the potential for commonly prescribed non-antibiotic medicines to undermine antibiotic therapy, warranting further study given the rising *S. aureus* treatment failures.

## Introduction

*Staphylococcus aureus* is a major human pathogen, responsible for over one million global deaths annually (1). Although commonly carried as a commensal, this opportunistic pathogen can cause a wide spectrum of infections, ranging from mild skin and soft-tissue infections to severe invasive infections, such as bacteraemia (2,3). Treatment options for these infections largely depend on the antibiotic susceptibility of the causative isolate. For methicillin-susceptible *S. aureus* (MSSA) infections, β-lactam antibiotics such as oxacillin are the treatment of choice. In contrast, therapeutic options for methicillin-resistant *S. aureus* (MRSA) are more limited, with recommended agents including vancomycin and daptomycin (4,5).

*S. aureus* infections are more common in elderly populations, with incidence increasing markedly with age and peaking in individuals over 80 years (6–8). Ageing is accompanied by an increased prevalence of comorbidities (9), which often necessitates the long-term use of multiple medications, and contributes to both increased susceptibility to infection and poorer clinical outcomes (10,11). Specifically, bacteraemia incidence is higher in this patient group and is associated with over 2-fold increase in mortality and is more likely to progress to complicated and chronic infection types such as osteomyelitis and infective endocarditis, compared to younger adults (12–16). Reflecting this greater burden of disease, patients with comorbidities receive a disproportionate number of antibiotic prescriptions (17). In the UK, the presence of comorbidities is associated with a 44% higher rate of antibiotic prescribing in primary care (18). Conditions most strongly linked to elevated antibiotic use include cardiovascular disease, diabetes, neurological disorders, and immunosuppression (19,20) This increased exposure to antibiotics, combined with frequent healthcare contact, blood tests, and invasive interventions, further elevates the risk of infection with drug-resistant pathogens in these populations (21). Indeed, elderly patients are also more likely to harbour MRSA infections, further complicating treatment and contributing to treatment failure (22).

Non-antibiotic medications prescribed to manage comorbidities have gained interest for potential repurposing, as some demonstrate off-target antimicrobial effects. For example, a large-scale systematic screen of human-targeted drugs against gut bacteria found that approximately 24% of these non-antibiotic drugs inhibited the growth of at least one bacterial species tested (23). While many studies have focused on understanding their antimicrobial mechanisms and potential synergy with antibiotics (24–27). It remains poorly understood whether these medications reduce antibiotic efficacy or select for resistance (21). Approximately 90% of people >65 take at least one regularly prescribed medicine, with a median of five per patient (20). Yet, the role of polypharmacy in causally contributing to the high burden of drug-resistant infections and associated morbidity and mortality remains unknown.

Here, we investigate the impact of eight non-antibiotic medicines, commonly prescribed for several indications associated with high antibiotic use (19,28), on antibiotic susceptibility of MRSA. This addresses a key gap in understanding the impact of commonly used medicines on resistance dynamics.

## Results

### Simvastatin, fluoxetine, and amlodipine exhibit anti-MRSA antimicrobial activity

Eight commonly prescribed non-antibiotic medicines_commonly prescribed for several indications, covering diverse drug classes, were chosen for their high incidence of use (28). For example, it is estimated that >7 million people in the UK take statins (29). A summary of these medicines, their indications, and their incidence of use is provided in Supplementary Table. 1. To determine whether these prescription non-antibiotic medicines exhibited antimicrobial activity, we monitored bacterial growth over 16 hours in the presence of a two-fold dilution series of each drug (Fig. 1a-h). Among the compounds tested, simvastatin, fluoxetine, and amlodipine showed notable activity against *S. aureus*, with minimum inhibitory concentrations (MICs) of 25, 50, and 100 µg/mL, respectively, which is in keeping with previous findings (Fig. 1a, b, c) (30–32). Although we were unable to determine an MIC for levothyroxine and omeprazole, we noted a dose-dependent delay in growth rate in the presence of these drugs (Fig. 1g, h). Together, this confirms that a subset of prescription non-antibiotic medicines possess antimicrobial activity against *S. aureus*.

**Figure 1.**
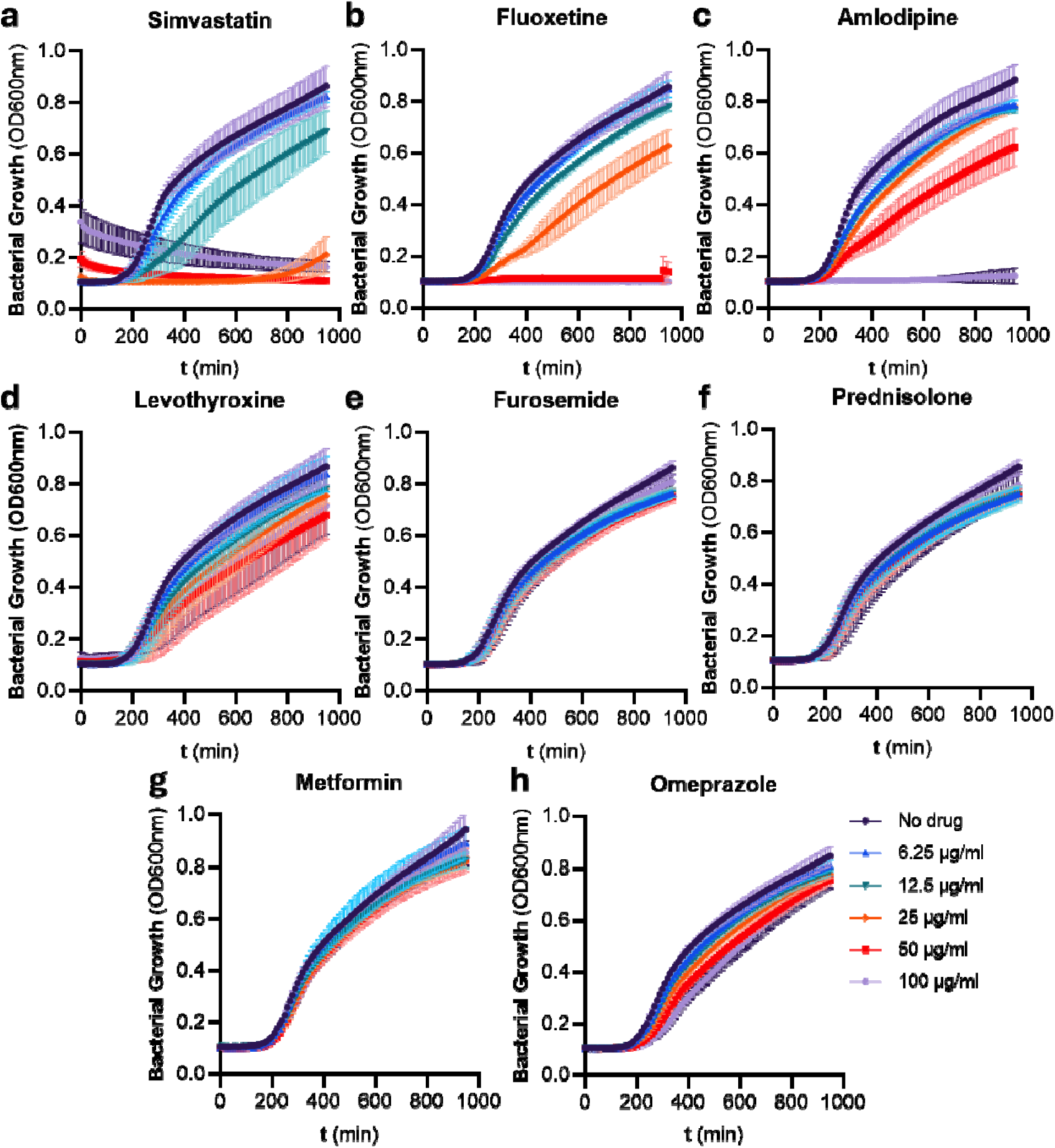
Simvastatin, amlodipine, and fluoxetine have antimicrobial activity. **(a-h)** 16-hour growth curves of *S. aureus* JE2 in the presence of a 1:2 dilution series of different prescription non-antibiotics. All experiments were replicated in n=3 independent assays. Error bars show the standard deviation of the mean. Note, the high initial starting optical density of simvastatin at 100 µg/ml is due to this concentration being close to the solubility threshold of this compound.

### Prescription non-antibiotics activate *S. aureus* stress responses

*S. aureus* has several dedicated signalling pathways, which are used to sense and respond to environmental stimuli such as the presence of antibiotics (33), particularly VraSR (cell wall damage) (34,35), GraSR (membrane damage) (36), and SOS (DNA damage) (37,38). Therefore, to better understand the antimicrobial activity of simvastatin, fluoxetine, and amlodipine, we tested whether they activated these key stress pathways. To achieve this, we used fluorescent reporter systems in which *gfp* expression was driven by the promoter of a gene activated by the relevant stress pathway. For the VraSR, GraSR, and SOS pathways, the promoters *vraX*, *dltA*, and *recA* were used, respectively, as they are well-established targets of activation by each pathway (39,40). A dose-dependent increase in *vraX* expression was observed in response to fluoxetine, amlodipine, and the positive control antibiotic vancomycin, confirming that these compounds induce cell wall stress (Fig. 2b, c. Supplementary Fig. 1a). However, activation of *vraX* expression was not detected in response to simvastatin (Fig. 2a). Similarly, *dltA* expression increased in a dose-dependent manner in response to fluoxetine, amlodipine, and the control antibiotic polymyxin B, but not simvastatin, indicating that the former compounds also induce membrane damage (Fig. 2a, b, c. Supplementary Fig. 1b). In contrast, neither of the three non-antibiotics activated *recA* expression, unlike the positive control antibiotic ciprofloxacin (Fig. 2a, b, c. Supplementary Fig. 1c).

**Figure 2.**
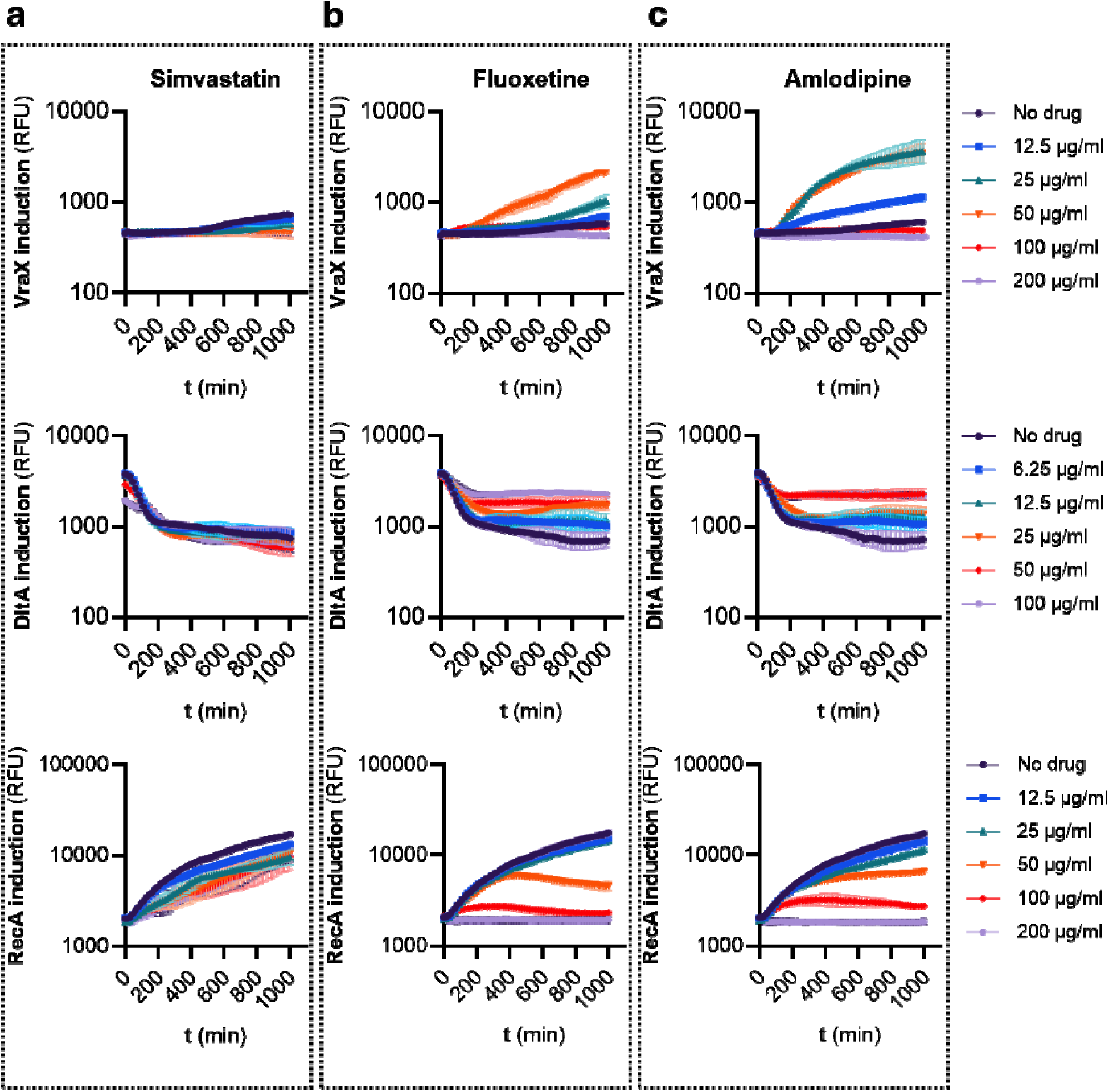
Fluoxetine and amlodipine activate the cell-wall and membrane stress responses. **(a)** Induction assays of p*vraX-*GFP, p*dltA-*GFP, and p*recA-*GFP in the presence of a 1:2 dilution series of simvastatin as determined by GFP accumulation over time. **(b)** Induction assays of p*vraX-*GFP, p*dltA-*GFP, p*recA-*GFP in the presence of a 1:2 dilution series of fluoxetine, as determined by GFP accumulation over time. **(c)** Induction assays of p*vraX-*GFP, p*dltA-*GFP, p*recA-*GFP in the presence of a 1:2 dilution series of amlodipine as determined by GFP accumulation over time. All experiments were replicated in n=3 independent assays in *S. aureus* JE2. Error bars show the standard deviation of the mean.

### Prescription non-antibiotic medicines damage the *S. aureus* membrane

Antimicrobials that simultaneously activate both cell wall and membrane stress responses are typically associated with a direct membrane-damaging mechanism of action, such as daptomycin (41–43). Therefore, to determine whether these prescription non-antibiotics also damage the bacterial membrane, we measured the ingress of the cell-impermeant nucleic acid stain Sytox Green in JE2 cells exposed to a 1:2 dilution series of amlodipine, fluoxetine, or simvastatin. Consistent with a direct membrane-damaging mechanism of action, both amlodipine and fluoxetine rapidly permeabilised the membrane, as evidenced by saturation of the Sytox fluorescence by 45 min (Fig. 3a, b). This rapid Sytox accumulation was associated with mild, dose-dependent cell lysis, further supporting the membrane-damaging activity of these drugs against *S. aureus* (Supplementary Fig. 2a, b). Both amlodipine and fluoxetine are cationic, with amlodipine bearing a primary amine and fluoxetine a secondary amine. Therefore, we hypothesised that the membrane damage caused by these drugs was mediated through electrostatic interactions with negatively charged phospholipids in the *S. aureus* membrane. Indeed, supplementation of the MHB with 10 µM phosphatidylglycerol increased the MICs of amlodipine and fluoxetine 4-fold.

**Figure 3.**
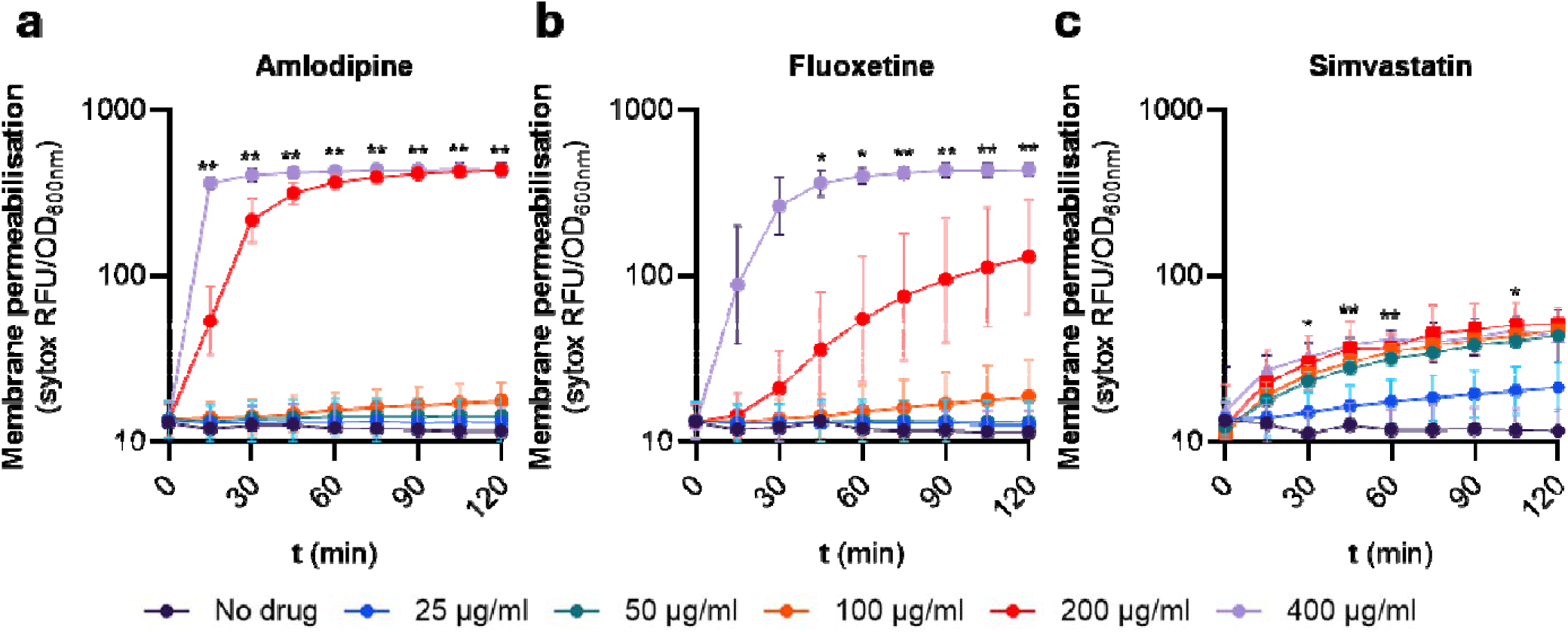
Amlodipine, fluoxetine and simvastatin permeabilise the membrane. **(a, b, c)** Membrane disruption of *S. aureus* JE2 exposed to a 1:2 dilution series of **(a)** amlodipine, **(b)** fluoxetine, **(c)** simvastatin, for 2 hours, as determined by uptake of the fluorescent nucleic acid dye Sytox green. RFU, relative fluorescence units. Significant differences were determined between the no drug condition and the 400 µg/ml treated condition by two-way repeated measures ANOVA with post hoc Dunnett’s test to correct for multiple comparisons. \**P□*<□0.05; \*\**P□*<□0.01.

Simvastatin exposure resulted in a slower and less pronounced accumulation of Sytox (Fig. 3b). This delayed kinetic profile is typical of antimicrobials that perturb phospholipid biosynthesis or other upstream biosynthetic processes, which indirectly compromise the membrane or cell envelope and weaken barrier function (44–46). Supporting this, proteomic and macromolecular-synthesis studies have shown that simvastatin inhibits multiple biosynthetic processes, including DNA, RNA, protein, cell-wall and lipid synthesis.(47).

### Transposon mutant screening reveals pleiotropic determinants of simvastatin resistance

Simvastatin inhibits HMG-CoA reductase in humans, blocking the rate-limiting step of cholesterol biosynthesis in the mevalonate pathway (48). In *S. aureus*, it concomitantly inhibits the bacterial HMG-CoA reductase homolog MvaA, disrupting isoprenoid, lipid, and peptidoglycan synthesis pathways essential for cell viability (Fig. 4a) (47,49–51). Further demonstrating simvastatin’s effect on the isoprenoid synthesis, we and others have shown that it inhibits the production of the isoprenoid-dependent carotenoid pigment staphyloxanthin (Fig. 4b) (52,53). Supplementation of exogenou mevalonate reversed the inhibitory activity of simvastatin on staphyloxanthin production, confirming it inhibitory activity on the isoprenoid synthesis pathway (Fig. 4c).

**Figure 4.**
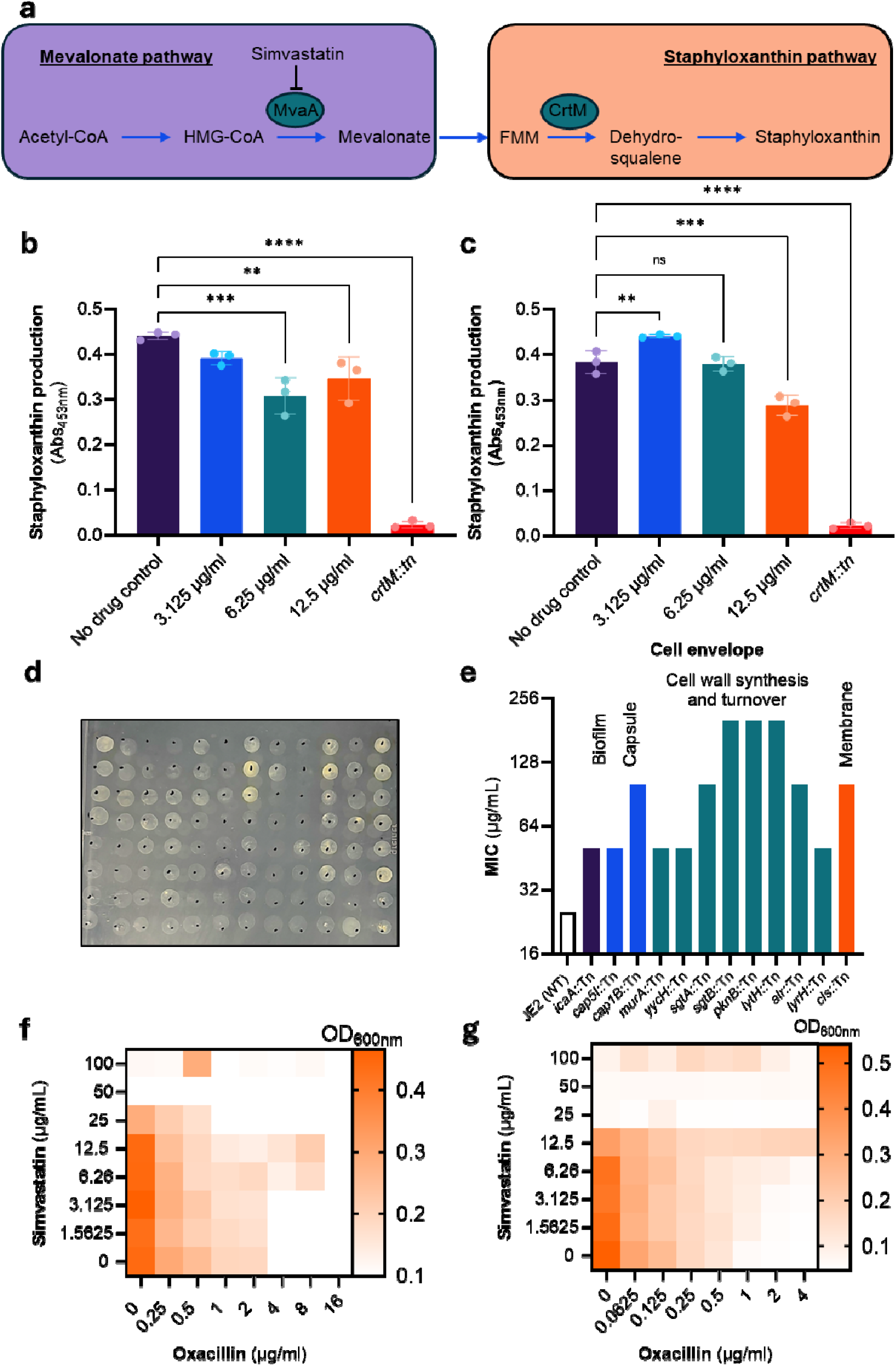
Screening of the NTML reveals pleiotropic routes for simvastatin resistance. **a)** Summary diagram of how the mevalonate pathway feeds into staphyloxanthin production. Acetyl-CoA is converted to 3-Hydroxy-3-methylglutaryl-CoA (HMG-CoA). This is subsequently converted to mevalonate by MvaA, which is inhibited by simvastatin. A multi-step reaction converts mevalonate into farnesyl pyrophosphate (FPP). FPP feeds into peptidoglycan, lipid and staphyloxanthin synthesis. CrtM catalyses the production of dehydrosqualene from FPP, ultimately resulting in the synthesis of staphyloxanthin. **b)** Staphyloxanthin abundance in *S. aureus* JE2 exposed, or not, to 3.125, 6.25, or 12.5 µg/mL simvastatin overnight. **c)** Staphyloxanthin abundance in *S. aureus* JE2 exposed, or not, to 3.125, 6.25, or 12.5 µg/mL simvastatin overnight, in the presence of exogenous mevalonate 10 µg/mL. JE2 *crtM::*Tn is used as a negative control. **d)** A representative plate from the NTML transposon screen. **e)** MIC of confirmed transposon mutant hits defective in cell envelope synthesis. **f)** Checkerboard broth microdilution assay showing the antagonistic interaction between simvastatin and oxacillin against *S. aureus* JE2. **g)** Checkerboard broth microdilution assay showing the antagonistic interaction between simvastatin and oxacillin against *S. aureus* Newman. Significant differences were determined by one-way repeated measures ANOVA between the no-drug control and simvastatin-treated conditions. \**P□*<□0.05; \*\**P□*<□0.01; \*\*\**P*<0.001; \*\*\*\**P*<0.0001; ns not significant.

To further elucidate simvastatin’s effects on *S. aureus* and identify genes and pathways that confer resistance, we screened the Nebraska Transposon Mutant Library (NTML) for mutants exhibiting reduced susceptibility at 1x MIC of simvastatin in Mueller-Hinton Broth agar (MHBA) (Fig. 4d)(54). The 101 mutants that grew at this concentration were tested for reduced simvastatin susceptibility by broth MIC testing, yielding 77 validated hits (Supplementary Table. 2). Of these hits, 12 were associated with cell envelope synthesis (Fig. 4e), 16 with DNA and protein synthesis (Supplementary Table 2), and 15 with metabolism and redox (Supplementary Table 2), highlighting the pleiotropic effect of inhibiting isoprenoid synthesis in *S. aureus.* The remaining hits (Supplementary Table. 2) were associated with virulence, transport, phage-associated genes, or had unknown function. Notably, several identified genes have previously been linked to altered susceptibility to cell wall antibiotics, including β-lactams (*murA*, *sgtA*, *sgtB*, *yycH*, *pknB*, *lytH*, *alr*, *clpC*) (55–61), vancomycin (*alr*, *yycH*, *sgtA*, *sgtB*, *clpC*) (56,59,62), and daptomycin (*cls*, *clpC*, *yycH*) (59,62,63). This overlap suggests that simvastatin’s inhibition of isoprenoid biosynthesis induces cell envelope damage analogous to these agents, potentially selecting for cross-resistance mutations while also creating opportunities for synergy.

Simvastatin has previously been shown to synergise with β-lactams against MRSA (64). Statins’ synergy arises from their disruption of fluid membrane microdomains (FMMs) enriched in isoprenoid lipids, thereby impairing PBP2a oligomerisation, which is essential for methicillin resistance. (64). To further test the potential for simvastatin to synergise with β-lactams, we performed a checkerboard broth microdilution assay (Fig. 4e). Unlike previously (64), we observed high levels of antagonism between oxacillin and simvastatin against JE2, with a fractional inhibitory concentration index (FICI) of >4 (Fig. 4f). In the presence of sub-inhibitory concentrations of simvastatin, the oxacillin MIC shifted 4-fold to 16 µg/ml (Fig. 4f). To test whether this phenotype extended to additional strains, we repeated the checkerboard analysis for the methicillin-susceptible *S. aureus* (MSSA) strain Newman (Fig. 4g). Similarly, simvastatin antagonised oxacillin susceptibility with a FICI >4. Furthermore, in the presence of 12.5 µg/ml, the oxacillin MIC shifted from 1 µg/ml (susceptible) to 4 µg/ml (resistant), according to CLSI breakpoints. Subsequent MIC testing of bacteria grown from these wells in the absence of simvastatin reverted the MIC to 1 µg/ml, potentially indicating that simvastatin may confer transient β-lactam resistance independent of PBP2A, more specifically, borderline oxacillin-resistant *S. aureus* (BORSA).

### Prescription non-antibiotic medicines accelerate resistance emergence to anti-MRSA antibiotics

First-line anti-MRSA antibiotics vancomycin and daptomycin activate the VraSR and GraSR stress responses, respectively. Having demonstrated that fluoxetine and amlodipine also activate the VraSR and GraSR stress pathways, and that simvastatin disrupts cell wall and membrane synthesis, we assessed whether these drugs could synergise with these antibiotics. Checkerboard broth microdilution assays were conducted to evaluate interactions between vancomycin and each of the following non-antibiotic agents: simvastatin, fluoxetine, and amlodipine (Fig. 5a, b, c). At sub-inhibitory concentrations, all three compounds reduced bacterial susceptibility to vancomycin, evidenced by an increase in the vancomycin MIC (Fig. 5a, b, c). Checkerboard broth microdilution assays were also conducted between daptomycin and these non-antibiotic agents (Fig. 5d, e, f). No evidence of synergy or antagonism was observed between daptomycin and simvastatin, or daptomycin and fluoxetine. However, sub-inhibitory concentrations of amlodipine antagonised daptomycin susceptibility, causing an increase in MIC from 1 µg/mL to 2 µg/mL (Fig. 5f). Collectively, these findings suggest that certain non-antibiotic drugs can impair antibiotic efficacy, potentially through shared mechanisms of action or the induction of cellular stress responses.

**Figure 5.**
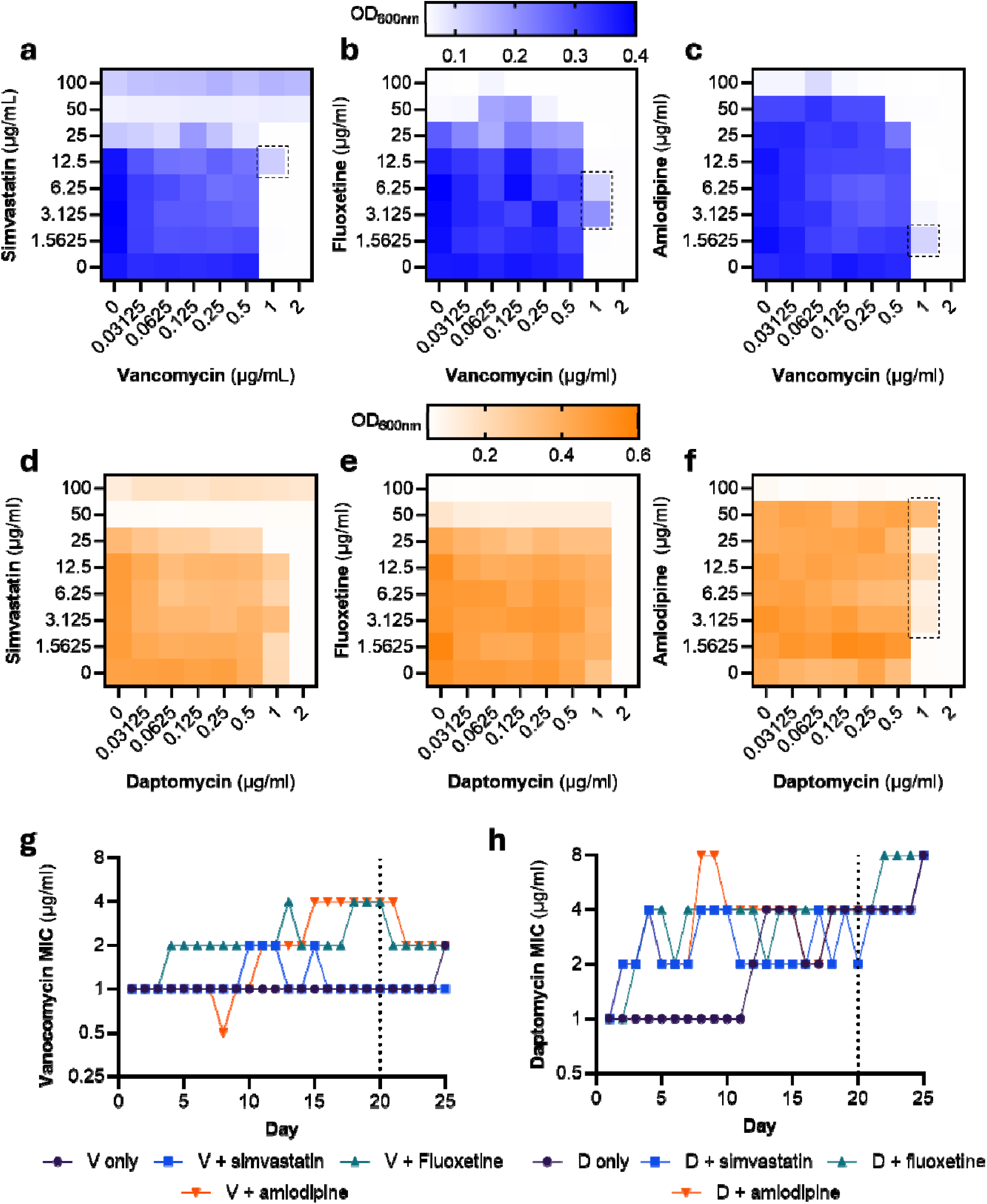
Simvastatin, fluoxetine, and amlodipine promote resistance to vancomycin and daptomycin. **(a, b, c)** Checkerboard broth microdilution assay showing the antagonistic interaction between vancomycin and **a)** simvastatin, **b)** fluoxetine, and **c)** amlodipine against *S. aureus* JE2. **(d, e, f)** Checkerboard broth microdilution assay between daptomycin and **a)** simvastatin, **b)** fluoxetine, **c)** amlodipine against *S. aureus* JE2. Cells displaying antagonism are highlighted by a dashed square. **g)** Increase of vancomycin MIC, following daily passage in either vancomycin (V) alone, or vancomycin in combination with subinhibitory simvastatin, fluoxetine, or amlodipine. **h)** Increase of vancomycin MIC, following daily passage in either daptomycin (D) alone, or daptomycin in combination with subinhibitory simvastatin, fluoxetine, or amlodipine. The dashed line at day 20 indicates where co-selection with simvastatin, fluoxetine, or amlodipine was removed.

Sustained activation of the VraSR and GraSR stress response systems has been linked to the development of resistance to vancomycin and daptomycin (65–69). Given the observed antagonism between the non-antibiotic agents and these antibiotics, together with evidence that the same compounds activate VraSR and GraSR, we hypothesised that these non-antibiotics may impose a selective pressure that promotes the emergence of vancomycin and daptomycin resistance. To test this, JE2 was passaged daily in vancomycin or daptomycin alone, or in combination with the non-antibiotic drugs (Fig. 5g, h). When passaged in vancomycin alone, the MIC shifted 2-fold on the 25^th^ passage (Fig. 5g). However, when vancomycin was combined with either simvastatin, fluoxetine, or amlodipine, MIC shifts were observed at earlier passage days (Fig. 5g). A 2-fold shift in vancomycin MIC was observed at day 4 for the vancomycin-fluoxetine combination, which rose to 4-fold (4 µg/mL) at day 18 (Fig. 5g). Similarly, the vancomycin MIC shifted 2-fold at day 11 for the vancomycin-amlodipine combination, further rising 4-fold at day 15 (Fig. 5g). Simvastatin also caused a 2-fold increase in vancomycin MIC; however, this was unstable at reverted to starting MIC (Fig. 5g). When non-antibiotic co-selection was removed at day 20, the vancomycin MIC decreased 2-fold, highlighting the selective pressure of these drugs.

Daptomycin serial passage resulted in a 2-fold increase in MIC at day 12, a 4-fold increase at day 18, and an 8-fold increase at day 25. In keeping with the antagonistic checkerboard analysis, co-selecting with simvastatin caused a 2-fold shift in MIC at day 2. The following passage days alternated between 2-fold and 4-fold increases in MIC, with a stable 4-fold increase in MIC achieved at day 21, and an 8-fold increase at day 25. A stable 4-fold increase in daptomycin MIC was also achieved at day 14, co-selecting with fluoxetine. An 8-fold increase in daptomycin MIC was also achieved earlier at day 22 for this combination. When passaged with amlodipine, the daptomycin MIC shifted sharply by 8-fold on day 8, followed by a 2-fold drop, which was maintained until day 25, where it again rose 2-fold to 8 µg/mL. Unlike the vancomycin passage, removal of the co-selecting non-antibiotic drugs did not influence the daptomycin MIC.

## Discussion

Older adults experience disproportionately high rates of drug-resistant infections, which are associated with poor clinical outcomes, including increased morbidity and mortality. Antibiotic use is also markedly elevated in this population, particularly among individuals with multiple comorbidities. Management of these chronic conditions often necessitates long-term pharmacotherapy, with the number of prescribed medications increasing substantially with age. Therefore, in this study, we asked whether this pattern of extensive medication use may contribute to antimicrobial resistance through complex drug interactions or poorly characterised additional selection pressures. We screened eight widely prescribed non-antibiotic medications, spanning therapeutic classes commonly used to manage comorbidities associated with high antibiotic use, for antimicrobial activity against MRSA. Interestingly, three were found to have robust antimicrobial activity: simvastatin, amlodipine, and fluoxetine. These three medications were selected for further characterisation of the antimicrobial mechanism of action and impact on antibiotic susceptibility, as we hypothesised that medications exhibiting off-target antimicrobial effects were more likely to modulate antibiotic efficacy.

We found that both amlodipine and fluoxetine activated cell wall and membrane stress responses, consistent with mechanisms involving disruption of membrane integrity and interference with cell wall synthesis. Membrane perturbation was further supported and demonstrated by increased accumulation of the fluorescent DNA stain Sytox Green. Beyond *S. aureus*, both agents have demonstrated antimicrobial activity against a range of bacterial species. However, the reduced potency observed in Gram-negative organisms suggests that these compounds likely act at the inner membrane, with activity constrained by the outer membrane permeability barrier that limits access to their target sites (70–72).

Although we observed antagonism between these agents and vancomycin, both drugs have been reported to synergise with other antibiotic classes in a context-dependent manner. For example, fluoxetine enhances the activity of erythromycin and gentamicin against *Pseudomonas aeruginosa*, *Escherichia coli*, and *S. aureus* (73), and has also been shown to potentiate meropenem, fosfomycin, and polymyxin B against a range of Gram-negative pathogens (74). This synergy has been attributed to inhibition of efflux, as fluoxetine exposure is associated with increased intracellular accumulation of ethidium bromide (75,76). Similarly, amlodipine has been shown to synergise with oxacillin against *S. aureus*, although this effect appears to be strain dependent (77). Nevertheless, the selection of fluoxetine-resistant mutants was associated with cross-resistance to multiple antibiotics, highlighting the potential for long-term exposure to these agents to undermine antibiotic efficacy (75). However, the accumulation of efflux-associated mutations is unlikely to confer resistance to a non-efflux substrate such as vancomycin. Therefore, the accelerated emergence of vancomycin resistance observed during co-selection with fluoxetine or amlodipine is likely independent of efflux-mediated mechanisms.

Removal of fluoxetine or amlodipine co-selection led to an almost immediate twofold reduction in vancomycin MIC. This is concerning, as the transient nature of this phenotype suggests that standard antibiotic susceptibility testing may fail to detect elevated vancomycin MICs in patients taking these medications. True vancomycin non-susceptibility is rare, with global prevalence estimates of 2.4% for vancomycin-resistant *S. aureus* (VRSA) and 4.3% for vancomycin-intermediate *S. aureus* (VISA) (78); by contrast, vancomycin treatment failure is relatively common, occurring in up to 50% of cases (79). Host-associated stressors that activate the VraSR and GraSR pathways have been shown to confer transient daptomycin and vancomycin tolerance (39), supporting the concept that environmental or physiological signals can temporarily modulate antibiotic susceptibility. It is therefore plausible that prescription medications such as fluoxetine or amlodipine may similarly contribute to vancomycin treatment failure, although the magnitude of this effect remains unknown. Simvastatin, fluoxetine, and amlodipine are among the most commonly prescribed medications in the UK, with millions of people receiving these drugs annually (29,80,81). Given this widespread use, the potential for these medications to influence antimicrobial susceptibility is almost certainly underappreciated.

While we did not detect activation of cell wall or membrane stress responses by simvastatin, previous studies have shown that it induces expression of *vraX*, other cell wall stress-associated genes, and *mvaA* of the mevalonate pathway (49). These discrepancies may reflect strain-specific regulation of cell wall stress response pathways (49). We were able to further verify the effect of simvastatin on disrupting cell wall integrity through a screen of the NTML, which yielded a number of hits responsible for cell wall synthesis or turnover. Notably, loss of *sgtB* and *murE* was associated with simvastatin resistance, consistent with previous reports showing their induction upon simvastatin exposure (49). We also observed a highly antagonistic interaction between simvastatin and oxacillin, which was able to confer transient BORSA in the MSSA strain Newman. This conflicts with previous reports of simvastatin β-lactam synergy (64). Interestingly, however, a large-scale screen for antibiotic-non-antibiotic drug interactions found that atorvastatin also antagonises β-lactams against *S. aureus* (82). Similarly, the antibiotic fosmidomycin, which acts independently of the mevalonate pathway but also depletes FPP levels, protects against lysis by the cell-wall antibiotic fosfomycin (83). Collectively, these findings indicate that statin–β-lactam interactions are highly context-dependent, likely influenced by strain background, pathway activity, and the β-lactam partner.

Polypharmacy (the concurrent use of multiple drugs) has been described as a silent pandemic, prompting initiatives focused on de-prescribing and reducing unnecessary medication burden to improve patient outcomes (84). Our findings suggest an additional, underappreciated benefit: targeted de-prescribing may help preserve antibiotic efficacy in already vulnerable populations, strengthening the rationale for accelerating these efforts. While such initiatives are clearly important, further research is needed to fully elucidate the impact of polypharmacy on antimicrobial resistance and to inform optimal strategies for medication optimisation.

## Methods

### Bacterial strains and growth conditions

The *S. aureus* strains used in this study are listed in Table 1. *S. aureus* was cultured in 3□ml Mueller-Hinton Broth (MHB) and grown overnight at 37□°C with shaking (180□rpm) to reach stationary phase. When necessary, MHB was supplemented with 10□µg/ml erythromycin or 90□µg/ml kanamycin. Since daptomycin requires calcium for activity, laboratory media were supplemented with 1.25 mM CaCl_2_. Prescription non-antibiotics used throughout this study are listed in Table 2. Unless stated otherwise, these were added to assays at a concentration of 12.5 µg/ml.

**Table 1.** Bacterial strains.

| Strain | Description | Reference |
| --- | --- | --- |
| USA300 JE2 (WT) | LAC strain of the USA300 CA-MRSA lineage cured of plasmids | (54) |
| USA300 JE2 *PvraX-gfp* | USA300 LAC JE2 carrying the *PvraX-gfp* reporter plasmid, Kan<sup>r</sup> | (39) |
| USA300 JE2 *PdltA-gfp* | USA300 LAC JE2 carrying the *PdltA-gfp* reporter plasmid, Kan<sup>r</sup> | (39) |
| USA300 JE2 *PrecA-gfp* | USA300 LAC JE2 carrying the *PrecA-gfp* reporter plasmid, Kan<sup>r</sup> | (40) |
| USA300 JE2 *crtM::Tn* | USA300 LAC JE2 with a *bursa aurealis* transposon insertion in *crtM*, Ery<sup>r</sup> | (54) |
| Newman | MSSA, laboratory strain isolated from human infection (CC8), lacks antibiotic-resistant determinants | (85) |

**Table 2.** Prescription non-antibiotic drugs.

| Non-antibiotic drugs | CAT number |
| --- | --- |
| Simvastatin | PHR1438 |
| Prednisolone | PHR1043 |
| Furosemide | PHR1057 |
| Fluoxetine hydrochloride | PHR1394 |
| Amlodipine besylate | PHR1185 |
| Levothyroxine | PHR1613 |
| Omeprazole | PHR1059 |
| Metformin | PHR1084 |

### Determination of MICs

The MIC of the prescription non-antibiotics were determined according to the well-established broth microdilution method (86). In brief, a 96-well microtitre plate was used to prepare a range of drug concentrations in 100□µl of cation-adjusted MHB via a series of 2-fold serial dilutions. To assess the impact of phospholipids on amlodipine and fluoxetine susceptibility, phosphatidylglycerol (Sigma-Aldrich) was added at a final concentration of 10 µM. For antibiotic synergy determination, checkerboard analyses were performed by preparing 2-fold serial dilutions of the drugs and antibiotics, with each dilution across a different axis, generating an 8□×□8 matrix to assess the MICs of each antibiotic when used in combination (87). Stationary-phase bacteria were diluted to 1□×□10^6^□c.f.u.□ml^−1^ in cation-adjusted MHB, and 100□µl was used to seed each well of the microtitre plate to give a final inoculation density of 5□×□10^5^□c.f.u.□ml^−1^. Plates were then incubated statically at 37□°C for 18□h in air, at which point the MIC was defined as the lowest antibiotic concentration at which no visible bacterial growth was observed. Where checkerboard analysis was performed, optical density (OD) was measured at 595_nm_ using a Bio-Rad iMark microplate absorbance reader (Bio-Rad Laboratories). Where growth curves are presented, plates were incubated at 37°C with continuous shaking in a microplate reader (Tecan Spark, MIC9412), and OD_600nm_ was recorded every 10 minutes for 16 hours.

### Fluorescent reporter assays

Promoter-*gfp* constructs were used to measure gene expression in response to amlodipine, simvastatin and fluoxetine, according to a pre-established protocol (39,40). In brief, amlodipine, simvastatin and fluoxetine were serially diluted 1:2 in 100 µl of cation-adjusted MHB (P*vraX-gfp,* P*recA-gfp*) or Rosewell Park Memorial Institute 1640 (RPMI) (Gibco, USA) (P*dltA-gfp*) in a black, clear-bottom 96-well plate (Greiner, Germany). Additionally, DMSO (Sigma-Aldrich) was added to all wells in the simvastatin dilution series at a final concentration of 2% to maintain drug solubility. Positive controls were run in parallel using vancomycin for *PvraX-gfp*, ciprofloxacin for *PrecA-gfp*, and polymyxin B for P*dltA-gfp*. Overnight reporter cultures were diluted to a final concentration of 10^8^ CFU/ml in MHB or RPMI 1640, and 100 μl of the diluted cultures was added to each well. The plates were placed in a microplate reader (Tecan Spark, MIC9412) at 37°C with shaking, and GFP fluorescence (excitation 475nm, emission 525nm) was measured every 15 minutes for 16 hours.

### Determination of membrane permeability

To measure membrane permeabilisation by amlodipine, simvastatin, and fluoxetine, the well-established SYTOX green assay was used (88,89). Stationary-phase *S. aureus,* at an inoculum density of 1□×□10^8^□c.f.u.□ml^−1^ was added to 3□ml of MHB containing the relevant prescription antibiotics across a 1:2 dilution series. The fluorescent probe SYTOX green (Invitrogen) was added to these cultures at a final concentration of 1□µM. Aliquots (200□μl) were transferred to a black microtitre plate with clear-bottomed wells (Greiner Bio-One). Fluorescence and OD_600nm_ were measured in a Tecan Infinite 200 Pro plate reader (excitation at 535□nm, emission at 617□nm) every 15□min for 2□h at 37□°C with shaking. Raw fluorescence readings were normalised according to OD_600nm_. OD_600nm_ measurements were also used to assess the extent of cell lysis following exposure to these drugs

### Screening of the NTML for simvastatin resistance determinants

The NTML comprises 1,920 individual mutants arrayed across five 384-well plates, which were used to screen for mutants conferring simvastatin resistance. Mutants from each of the five plates were used to inoculate twenty 96-well microtiter plates, with each well containing MHB supplemented with 10 μg/mL of erythromycin (Sigma-Aldrich) to maintain selection for the transposon insertion. These 96-well plates were incubated statically at 37°C overnight. For storage, the NTML library was stored in 25% glycerol at - 80°C. For the screening assay, square agar plates containing MHB were prepared with simvastatin at 1x simvastatin MIC (25 μg/mL). After overnight growth, cultures from each well of the 96-well plates were spotted (3.5 μl) on the simvastatin agar plates. Plates were again statically incubated overnight at 37°C. Mutant identities were determined by cross-referencing the NTML plate map.

### Staphyloxanthin pigment analysis

A 3 mL overnight culture of *S. aureus* JE2 and the control strain JE2 *crtM*::Tn was prepared in MHB. The *S. aureus* JE2 MHB culture was supplemented with varying concentrations of simvastatin (0 μg/mL, 3.125 μg/mL, 6.25 μg/mL, and 12.5 μg/mL), with and without the addition of mevalonate (90469, Sigma Aldrich) at a concentration of 100 μM. The following day, 1 ml of the overnight cultures was centrifuged at 10,000 x g for 1 minute. The pellets were then resuspended in 200 μl of methanol and incubated at 55°C in a water bath for 30 minutes. Following incubation, the pellets were centrifuged at 10,000 x g for 1 minute. The supernatants were recovered, and the absorbance was read at 453 nm (Tecan Spark, MIC9412). A methanol-only control was included as a blank.

### Serial passaging with antibiotics

A single colony of *S. aureus* JE2 was grown overnight in 3 mL MHB. Two 96-well plates were prepared, one for vancomycin and one for daptomycin passaging. In brief, a 96-well microtitre plate was used to prepare a range of daptomycin and vancomycin concentrations in 100□µl of cation-adjusted MHB via a series of 2-fold serial dilutions. Following this, stationary-phase bacteria were diluted to 1□×□10^6^□c.f.u.□ml^−1^ in cation-adjusted MHB, with and without simvastatin, amlodipine, or fluoxetine at a final concentration of 12.5 μg/ml, and 100□µl used to seed each well of the microtitre plate to give a final inoculation density of 5□×□10^5^□c.f.u.□ml^−1^. Plates were then incubated statically at 37□°C for 18□h in air. Each day, bacteria from the well immediately adjacent to the MIC (the highest concentration in which there was observable growth) were diluted 1:5000 in fresh MHB and plated on a new microtitre prepared as above. Serial passaging was performed for 20 days, with daily OD_600nm_ measurements (Tecan Spark, MIC9412) used to monitor changes in MIC in the presence of each drug. This was followed by an additional 5-day passage in the absence of simvastatin, amlodipine, and fluoxetine to assess the stability of the resistance phenotype.

### Statistical analysis

The data is derived from three biologically independent experiments unless stated otherwise. The data is presented as the mean of the biological repeats with error bars representing the standard deviation from the mean. Where appropriate, statistical analysis is performed using an ordinary one-way or two-way analysis of variance (ANOVA) as stated. Details of the post hoc comparison and the asterisks representing significance levels are stated in the figure legends. All statistical analyses were performed using GraphPad Prism software Version 10.5.0.

## Acknowledgements

All authors acknowledge the provision of strains by the Network on Antimicrobial Resistance in *Staphylococcus aureus* (NARSA) Program under NIAID/ NIH Contract No. HHSN272200700055C. The funders had no role in the study design, interpretation of the findings or the writing of the manuscript.

## References

1. Naghavi M, Mestrovic T, Gray A, Gershberg Hayoon A, Swetschinski LR, Robles Aguilar G, et al. Global burden associated with 85 pathogens in 2019: a systematic analysis for the Global Burden of Disease Study 2019. Lancet Infect Dis. 2024 Apr. doi:10.1016/S1473-3099(24)00158-0

2. Lowy FD. Staphylococcus aureus Infections. New England Journal of Medicine. 1998 Aug 20;339(8):520–32. doi:10.1056/NEJM199808203390806

3. Tong SYC, Fowler VG, Skalla L, Holland TL. Management of Staphylococcus aureus Bacteremia. JAMA. 2025 Sep 2;334(9):798. doi:10.1001/jama.2025.4288

4. Nathwani D, Morgan M, Masterton RG, Dryden M, Cookson BD, French G, et al. Guidelines for UK practice for the diagnosis and management of methicillin-resistant Staphylococcus aureus (MRSA) infections presenting in the community. Journal of Antimicrobial Chemotherapy. 2008 May;61(5):976–94. doi:10.1093/jac/dkn096

5. Gordon RJ, Lowy FD. Pathogenesis of Methicillin-Resistant Staphylococcus aureus Infection. Clinical Infectious Diseases. 2008 Jun;46(S5):S350–9. doi:10.1086/533591

6. Thorlacius-Ussing L, Sandholdt H, Larsen AR, Petersen A, Benfield T. Age-Dependent Increase in Incidence of Staphylococcus aureus Bacteremia, Denmark, 2008–2015. Emerg Infect Dis. 2019 May;25(5). doi:10.3201/eid2505.181733

7. Tacconelli E, Pop-Vicas AE, D’Agata EMC. Increased mortality among elderly patients with meticillin-resistant Staphylococcus aureus bacteraemia. Journal of Hospital Infection. 2006 Nov;64(3):251–6. doi:10.1016/j.jhin.2006.07.001

8. Kang CI, Song JH, Ko KS, Chung DR, Peck KR. Clinical features and outcome of Staphylococcus aureus infection in elderly versus younger adult patients. International Journal of Infectious Diseases. 2011 Jan;15(1):e58–62. doi:10.1016/j.ijid.2010.09.012

9. Valderas JM, Starfield B, Sibbald B, Salisbury C, Roland M. Defining Comorbidity: Implications for Understanding Health and Health Services. The Annals of Family Medicine. 2009 Jul 1;7(4):357–63. doi:10.1370/afm.983

10. Lukovac E, KoluderCimic N, HadzovicCengic M, Baljic R, Hadzic A, Gojak R. Analysis of Comorbidity of the Patients Affected by Staphylococcal Bacteremia/Sepsis in the Last Ten Years. Materia Socio Medica. 2012;24(1):13. doi:10.5455/msm.2012.24.s13-s15

11. Valobdás NB, Alves MR, Silva EA dos SR da, Lourenço MC da S, Nascimento BC de N, Veloso VG, et al. Severe Staphylococcus aureus infection: associated factors and outcomes. The Brazilian Journal of Infectious Diseases. 2025 Sep;29(5):104573. doi:10.1016/j.bjid.2025.104573

12. Thorlacius-Ussing L, Sandholdt H, Larsen AR, Petersen A, Benfield T. Age-Dependent Increase in Incidence of Staphylococcus aureus Bacteremia, Denmark, 2008-2015. Emerg Infect Dis. 2019 May;25(5):875–82. doi:10.3201/eid2505.181733 PubMed PMID: 31002300.

13. Yahav D, Schlesinger A, Shaked H, Goldberg E, Paul M, Bishara J, et al. Clinical presentation, management and outcomes of Staph aureus bacteremia (SAB) in older adults. Aging Clin Exp Res. 2017 Apr 12;29(2):127–33. doi:10.1007/s40520-016-0543-4

14. Morimoto T, Hirata H, Otani K, Nakamura E, Miyakoshi N, Terashima Y, et al. Vertebral Osteomyelitis and Infective Endocarditis Co-Infection. J Clin Med. 2022 Apr 18;11(8):2266. doi:10.3390/jcm11082266

15. Douiyeb S, Sigaloff KCE, Ulas EG, Duffels MGJ, Drexhage O, Germans T, et al. Vertebral osteomyelitis in patients with infective endocarditis: prevalence, risk factors and mortality. European Journal of Clinical Microbiology & Infectious Diseases. 2025 Apr 21;44(4):819–25. doi:10.1007/s10096-025-05041-8

16. Ohta R, Sano C. Factors Affecting Recurrent Staphylococcus aureus Bacteremia Among Older Patients in Rural Community Hospitals: A Retrospective Cohort Study. Cureus. 2024 Sep;16(9):e70120. doi:10.7759/cureus.70120 PubMed PMID: 39449886.

17. Seo H, Sim YS, Min KH, Lee JH, Kim BK, Oh YM, et al. The Relationship Between Comorbidities and Microbiologic Findings in Patients with Acute Exacerbation of Chronic Obstructive Pulmonary Disease. Int J Chron Obstruct Pulmon Dis. 2022 Apr;Volume 17:855–67. doi:10.2147/COPD.S360222

18. Shallcross L, Beckley N, Rait G, Hayward A, Petersen I. Antibiotic prescribing frequency amongst patients in primary care: a cohort study using electronic health records. Journal of Antimicrobial Chemotherapy. 2017 Jun;72(6):1818–24. doi:10.1093/jac/dkx048

19. Fernández-Urrusuno R, Meseguer Barros CM, Anaya-Ordóñez S, Borrego Izquierdo Y, Lallana-Álvarez MJ, Madridejos R, et al. Patients receiving a high burden of antibiotics in the community in Spain: a cross-sectional study. Pharmacol Res Perspect. 2021 Feb 19;9(1). doi:10.1002/prp2.692

20. Christensen LD, Reilev M, Juul-Larsen HG, Jørgensen LM, Kaae S, Andersen O, et al. Use of prescription drugs in the older adult population—a nationwide pharmacoepidemiological study. Eur J Clin Pharmacol. 2019 Aug 4;75(8):1125–33. doi:10.1007/s00228-019-02669-2

21. Rockenschaub P, Hayward A, Shallcross L. Antibiotic Prescribing Before and After the Diagnosis of Comorbidity: A Cohort Study Using Primary Care Electronic Health Records. Clinical Infectious Diseases. 2020 Oct 23;71(7):e50–7. doi:10.1093/cid/ciz1016

22. Xi L, Li S, Chen M, Huang X, Li N, Chen N, et al. Age-Related Differences in Vancomycin-Associated Nephrotoxicity and Efficacy in Methicillin-Resistant Staphylococcus aureus Infection: A Comparative Study between Elderly and Adult Patients. Antibiotics. 2024 Apr 3;13(4):324. doi:10.3390/antibiotics13040324

23. Maier L, Pruteanu M, Kuhn M, Zeller G, Telzerow A, Anderson EE, et al. Extensive impact of non-antibiotic drugs on human gut bacteria. Nature. 2018 Mar 29;555(7698):623–8. doi:10.1038/nature25979

24. Younis W, Thangamani S, Seleem M. Repurposing Non-Antimicrobial Drugs and Clinical Molecules to Treat Bacterial Infections. Curr Pharm Des. 2015 Sep 22;21(28):4106–11. doi:10.2174/1381612821666150506154434

25. Tiwana G, Cock IE, Taylor SM, Cheesman MJ. Beyond Antibiotics: Repurposing Non-Antibiotic Drugs as Novel Antibacterial Agents to Combat Resistance. Int J Mol Sci. 2025 Oct 10;26(20):9880. doi:10.3390/ijms26209880

26. Dalhoff A. Are antibacterial effects of non-antibiotic drugs random or purposeful because of a common evolutionary origin of bacterial and mammalian targets? Infection. 2021 Aug 15;49(4):569–89. doi:10.1007/s15010-020-01547-9

27. Ejim L, Farha MA, Falconer SB, Wildenhain J, Coombes BK, Tyers M, et al. Combinations of antibiotics and nonantibiotic drugs enhance antimicrobial efficacy. Nat Chem Biol. 2011 Jun 24;7(6):348–50. doi:10.1038/nchembio.559

28. Audi S, Burrage DR, Lonsdale DO, Pontefract S, Coleman JJ, Hitchings AW, et al. The ‘top 100’ drugs and classes in England: an updated ‘starter formulary’ for trainee prescribers. Br J Clin Pharmacol. 2018 Nov 10;84(11):2562–71. doi:10.1111/bcp.13709

29. Trusler D. Statin prescriptions in UK now total a million each week. BMJ. 2011 Jul 14;343(jul14 2):d4350–d4350. doi:10.1136/bmj.d4350

30. Ahmed SA, Jordan RL, Isseroff RR, Lenhard JR. Potential Synergy of Fluoxetine and Antibacterial Agents Against Skin and Soft Tissue Pathogens and Drug-Resistant Organisms. Antibiotics. 2024 Dec 3;13(12):1165. doi:10.3390/antibiotics13121165

31. Boyd NK, Lee GC, Teng C, Frei CR. In vitro activity of non-antibiotic drugs against Staphylococcus aureus clinical strains. J Glob Antimicrob Resist. 2021 Dec;27:167–71. doi:10.1016/j.jgar.2021.09.003

32. Graziano TS, Cuzzullin MC, Franco GC, Schwartz-Filho HO, de Andrade ED, Groppo FC, et al. Statins and Antimicrobial Effects: Simvastatin as a Potential Drug against Staphylococcus aureus Biofilm. PLoS One. 2015 May 28;10(5):e0128098. doi:10.1371/journal.pone.0128098

33. Bleul L, Francois P, Wolz C. Two-Component Systems of S. aureus: Signaling and Sensing Mechanisms. Genes (Basel). 2021 Dec 23;13(1):34. doi:10.3390/genes13010034

34. Yin S, Daum RS, Boyle-Vavra S. VraSR two-component regulatory system and its role in induction of pbp2 and vraSR expression by cell wall antimicrobials in Staphylococcus aureus. Antimicrob Agents Chemother. 2006 Jan;50(1):336–43. doi:10.1128/AAC.50.1.336-343.2006 PubMed PMID: 16377706.

35. Gardete S, Wu SW, Gill S, Tomasz A. Role of VraSR in Antibiotic Resistance and Antibiotic-Induced Stress Response in Staphylococcus aureus. Antimicrob Agents Chemother. 2006 Oct;50(10):3424–34. doi:10.1128/AAC.00356-06

36. Yang SJ, Bayer AS, Mishra NN, Meehl M, Ledala N, Yeaman MR, et al. The Staphylococcus aureus two-component regulatory system, GraRS, senses and confers resistance to selected cationic antimicrobial peptides. Infect Immun. 2012 Jan;80(1):74–81. doi:10.1128/IAI.05669-11 PubMed PMID: 21986630.

37. Clarke RS, Ha KP, Edwards AM. RexAB Promotes the Survival of Staphylococcus aureus Exposed to Multiple Classes of Antibiotics. Antimicrob Agents Chemother. 2021 Sep 17;65(10). doi:10.1128/AAC.00594-21

38. Cheng K, Sun Y, Yu H, Hu Y, He Y, Shen Y. Staphylococcus aureus SOS response: Activation, impact, and drug targets. mLife. 2024 Sep;3(3):343–66. doi:10.1002/mlf2.12137 PubMed PMID: 39359682.

39. Ledger EVK, Mesnage S, Edwards AM. Human serum triggers antibiotic tolerance in Staphylococcus aureus. Nat Commun. 2022 Apr 19;13(1):2041. doi:10.1038/s41467-022-29717-3

40. Clarke RS, Bruderer MS, Ha KP, Edwards AM. RexAB Is Essential for the Mutagenic Repair of Staphylococcus aureus DNA Damage Caused by Co-trimoxazole. Antimicrob Agents Chemother. 2019 Dec;63(12). doi:10.1128/AAC.00944-19

41. Buttress JA, Schäfer AB, Koh A, Wheatley J, Mickiewicz K, Wenzel M, et al. The last resort antibiotic daptomycin exhibits two independent antibacterial mechanisms of action. Nat Commun. 2025 Nov 24;16(1):10320. doi:10.1038/s41467-025-65287-w

42. Müller A, Wenzel M, Strahl H, Grein F, Saaki TN V., Kohl B, et al. Daptomycin inhibits cell envelope synthesis by interfering with fluid membrane microdomains. Proceedings of the National Academy of Sciences. 2016 Nov 8;113(45). doi:10.1073/pnas.1611173113

43. Muthaiyan A, Silverman JA, Jayaswal RK, Wilkinson BJ. Transcriptional Profiling Reveals that Daptomycin Induces the Staphylococcus aureus Cell Wall Stress Stimulon and Genes Responsive to Membrane Depolarization. Antimicrob Agents Chemother. 2008 Mar;52(3):980–90. doi:10.1128/AAC.01121-07

44. Chamberlin J, Story S, Ranjan N, Chesser G, Arya DP. Gram-negative synergy and mechanism of action of alkynyl bisbenzimidazoles. Sci Rep. 2019 Oct 2;9(1):14171. doi:10.1038/s41598-019-48898-4

45. Pérez-Peinado C, Dias SA, Domingues MM, Benfield AH, Freire JM, Rádis-Baptista G, et al. Mechanisms of bacterial membrane permeabilization by crotalicidin (Ctn) and its fragment Ctn(15–34), antimicrobial peptides from rattlesnake venom. Journal of Biological Chemistry. 2018 Feb;293(5):1536–49. doi:10.1074/jbc.RA117.000125

46. Podoll J, Olson J, Wang W, Wang X. A Cell-Free Screen for Bacterial Membrane Disruptors Identifies Mefloquine as a Novel Antibiotic Adjuvant. Antibiotics. 2021 Mar 18;10(3):315. doi:10.3390/antibiotics10030315

47. Thangamani S, Mohammad H, Abushahba MFN, Hamed MI, Sobreira TJP, Hedrick VE, et al. Exploring simvastatin, an antihyperlipidemic drug, as a potential topical antibacterial agent. Sci Rep. 2015 Nov 10;5(1):16407. doi:10.1038/srep16407

48. Taylor F, Huffman MD, Macedo AF, Moore TH, Burke M, Davey Smith G, et al. Statins for the primary prevention of cardiovascular disease. Cochrane Database of Systematic Reviews. 2013 Jan 31;2021(9). doi:10.1002/14651858.CD004816.pub5

49. Cortês IT, Silva K de P, Cogo-Müller K. Effects of simvastatin on the mevalonate pathway and cell wall integrity of Staphylococcus aureus. J Appl Microbiol. 2025 Jan 6;136(1). doi:10.1093/jambio/lxaf012

50. Matsumoto Y, Yasukawa J, Ishii M, Hayashi Y, Miyazaki S, Sekimizu K. A critical role of mevalonate for peptidoglycan synthesis in Staphylococcus aureus. Sci Rep. 2016 Mar 10;6(1):22894. doi:10.1038/srep22894

51. Balibar CJ, Shen X, Tao J. The Mevalonate Pathway of Staphylococcus aureus. J Bacteriol. 2009 Feb;191(3):851–61. doi:10.1128/JB.01357-08

52. Chen G, Fan Z, Jia M, Li R, He Y, Lin X, et al. Simvastatin Administration Prohibits Staphylococcus aureus Anti-ROS Adaptation In Vivo and Alleviates Bone Infections. ACS Infect Dis. 2025 Sep 12;11(9):2466–75. doi:10.1021/acsinfecdis.5c00309

53. Burtchett TA, Ottosen EN, Jitsukawa T, Kaneko M, Yasui M, Lysne JA, et al. A redundant isoprenoid biosynthetic pathway supports Staphylococcus aureus metabolic versatility. mBio. 2025 Aug 13;16(8). doi:10.1128/mbio.00353-25

54. Fey PD, Endres JL, Yajjala VK, Widhelm TJ, Boissy RJ, Bose JL, et al. A Genetic Resource for Rapid and Comprehensive Phenotype Screening of Nonessential Staphylococcus aureus Genes. mBio. 2013 Mar;4(1). doi:10.1128/mBio.00537-12

55. Bæk KT, Gründling A, Mogensen RG, Thøgersen L, Petersen A, Paulander W, et al. β-Lactam Resistance in Methicillin-Resistant Staphylococcus aureus USA300 Is Increased by Inactivation of the ClpXP Protease. Antimicrob Agents Chemother. 2014 Aug;58(8):4593–603. doi:10.1128/AAC.02802-14

56. Suzuki Y, Kawada-Matsuo M, Thuan VTT, Le MNT, Sakaguchi T, Komatsuzawa H. D-alanine synthesis and exogenous alanine affect the antimicrobial susceptibility of Staphylococcus aureus. Antimicrob Agents Chemother. 2025 Jun 12. doi:10.1128/aac.01936-24

57. Fujimura T, Murakami K. Staphylococcus aureus Clinical Isolate with High-Level Methicillin Resistance with an lytH Mutation Caused by IS 1182 Insertion. Antimicrob Agents Chemother. 2008 Feb;52(2):643–7. doi:10.1128/AAC.00395-07

58. Huemer M, Mairpady Shambat S, Hertegonne S, Bergada-Pijuan J, Chang CC, Pereira S, et al. Serine-threonine phosphoregulation by PknB and Stp contributes to quiescence and antibiotic tolerance in Staphylococcus aureus. Sci Signal. 2023 Jan 3;16(766). doi:10.1126/scisignal.abj8194

59. Bleul L, Francois P, Wolz C. Two-Component Systems of S. aureus: Signaling and Sensing Mechanisms. Genes (Basel). 2021 Dec 23;13(1):34. doi:10.3390/genes13010034

60. Karinou E, Schuster CF, Pazos M, Vollmer W, Gründling A. Inactivation of the Monofunctional Peptidoglycan Glycosyltransferase SgtB Allows Staphylococcus aureus To Survive in the Absence of Lipoteichoic Acid. J Bacteriol. 2019 Jan;201(1). doi:10.1128/JB.00574-18

61. Gardete S, Ludovice AM, Sobral RG, Filipe SR, de Lencastre H, Tomasz A. Role of murE in the Expression of β-Lactam Antibiotic Resistance in Staphylococcus aureus. J Bacteriol. 2004 Mar 15;186(6):1705–13. doi:10.1128/JB.186.6.1705-1713.2004

62. Xu L, Henriksen C, Mebus V, Guérillot R, Petersen A, Jacques N, et al. A Clinically Selected Staphylococcus aureus clpP Mutant Survives Daptomycin Treatment by Reducing Binding of the Antibiotic and Adapting a Rod-Shaped Morphology. Antimicrob Agents Chemother. 2023 Jun 15;67(6). doi:10.1128/aac.00328-23

63. Lim C, Coombs GW, Mowlaboccus S. Molecular mechanisms of resistance and tolerance of Staphylococcus aureus to daptomycin. Int J Antimicrob Agents. 2026 Jan;67(1):107678. doi:10.1016/j.ijantimicag.2025.107678

64. García-Fernández E, Koch G, Wagner RM, Fekete A, Stengel ST, Schneider J, et al. Membrane Microdomain Disassembly Inhibits MRSA Antibiotic Resistance. Cell. 2017 Nov;171(6):1354–1367.e20. doi:10.1016/j.cell.2017.10.012

65. Taglialegna A, Varela MC, Rosato RR, Rosato AE. VraSR and Virulence Trait Modulation during Daptomycin Resistance in Methicillin-Resistant Staphylococcus aureus Infection. mSphere. 2019 Feb 27;4(1). doi:10.1128/mSphere.00557-18

66. Sabat AJ, Tinelli M, Grundmann H, Akkerboom V, Monaco M, Del Grosso M, et al. Daptomycin Resistant Staphylococcus aureus Clinical Strain With Novel Non-synonymous Mutations in the mprF and vraS Genes: A New Insight Into Daptomycin Resistance. Front Microbiol. 2018;9:2705. doi:10.3389/fmicb.2018.02705 PubMed PMID: 30459746.

67. Doddangoudar VC, Boost MV, Tsang DNC, O’Donoghue MM. Tracking changes in the vraSR and graSR two component regulatory systems during the development and loss of vancomycin non-susceptibility in a clinical isolate. Clinical Microbiology and Infection. 2011 Aug;17(8):1268–72. doi:10.1111/j.1469-0691.2011.03463.x

68. Yoo J Il, Kim JW, Kang GS, Kim HS, Yoo JS, Lee YS. Prevalence of amino acid changes in the yvqF, vraSR, graSR, and tcaRAB genes from vancomycin intermediate resistant Staphylococcus aureus. Journal of Microbiology. 2013 Apr 27;51(2):160–5. doi:10.1007/s12275-013-3088-7

69. Cui L, Neoh H min, Shoji M, Hiramatsu K. Contribution of vraSR and graSR Point Mutations to Vancomycin Resistance in Vancomycin-Intermediate Staphylococcus aureus. Antimicrob Agents Chemother. 2009 Mar;53(3):1231–4. doi:10.1128/AAC.01173-08

70. Rukavishnikov G, Leonova L, Kasyanov E, Leonov V, Neznanov N, Mazo G. Antimicrobial activity of antidepressants on normal gut microbiota: Results of the in vitro study. Front Behav Neurosci. 2023 Mar 22;17. doi:10.3389/fnbeh.2023.1132127

71. Sharma P, Kalra A, Tripathi AD, Chaturvedi VK, Chouhan B. Antimicrobial Proficiency of Amlodipine: Investigating its Impact on Pseudomonas spp. in Urinary Tract Infections. Indian J Microbiol. 2025 Mar 18;65(1):347–58. doi:10.1007/s12088-024-01280-z

72. Kumar KA, Ganguly K, Mazumdar K, Dutta NK, Dastidar SG, Chakrabarty AN. Amlodipine: a cardiovascular drug with powerful antimicrobial property. Acta Microbiol Pol. 2003;52(3):285–92. PubMed PMID: 14743981.

73. Karine de Sousa A, Rocha JE, Gonçalves de Souza T, Sampaio de Freitas T, Ribeiro-Filho J, Melo Coutinho HD. New roles of fluoxetine in pharmacology: Antibacterial effect and modulation of antibiotic activity. Microb Pathog. 2018 Oct;123:368–71. doi:10.1016/j.micpath.2018.07.040

74. Ahmed SA, Jordan RL, Isseroff RR, Lenhard JR. Potential Synergy of Fluoxetine and Antibacterial Agents Against Skin and Soft Tissue Pathogens and Drug-Resistant Organisms. Antibiotics. 2024 Dec 3;13(12):1165. doi:10.3390/antibiotics13121165

75. Jin M, Lu J, Chen Z, Nguyen SH, Mao L, Li J, et al. Antidepressant fluoxetine induces multiple antibiotics resistance in Escherichia coli via ROS-mediated mutagenesis. Environ Int. 2018 Nov;120:421–30. doi:10.1016/j.envint.2018.07.046

76. Nzakizwanayo J, Scavone P, Jamshidi S, Hawthorne JA, Pelling H, Dedi C, et al. Fluoxetine and thioridazine inhibit efflux and attenuate crystalline biofilm formation by Proteus mirabilis. Sci Rep. 2017 Sep 22;7(1):12222. doi:10.1038/s41598-017-12445-w

77. Barbosa AD, Sá LG, Neto JB, Rodrigues DS, Cabral VP, Moreira LE, et al. Activity of Amlodipine Against Staphylococcus Aureus1: Association with Oxacillin and Mechanism of Action. Future Microbiol. 2023 May 19;18(8):505–19. doi:10.2217/fmb-2022-0230

78. Shariati A, Dadashi M, Moghadam MT, van Belkum A, Yaslianifard S, Darban-Sarokhalil D. Global prevalence and distribution of vancomycin resistant, vancomycin intermediate and heterogeneously vancomycin intermediate Staphylococcus aureus clinical isolates: a systematic review and meta-analysis. Sci Rep. 2020 Jul 29;10(1):12689. doi:10.1038/s41598-020-69058-z

79. Kollef MH. Limitations of Vancomycin in the Management of Resistant Staphylococcal Infections. Clinical Infectious Diseases. 2007 Sep 15;45(Supplement_3):S191–5. doi:10.1086/519470

80. NHSBSA. NHS Business Services Authority. Prescription Cost Analysis - England 2024/25. 2025.

81. Jones M, Krockow EM, Tromans SJ, Mukaetova-Ladinska EB. Antidepressant prescribing trends for adult patients in the UK and Ireland during the COVID-19 pandemic: systematic review. BJPsych Open. 2026 Mar 2;12(2):e77. doi:10.1192/bjo.2026.10990

82. Cacace E, Kim V, Varik V, Knopp M, Tietgen M, Brauer-Nikonow A, et al. Systematic analysis of drug combinations against Gram-positive bacteria. Nat Microbiol. 2023 Sep 28;8(11):2196–212. doi:10.1038/s41564-023-01486-9

83. Kawai Y, Errington J. Antibiotic fosmidomycin protects bacteria from cell wall perturbations by antagonizing oxidative damage-mediated cell lysis. Front Microbiol. 2025 Apr 16;16. doi:10.3389/fmicb.2025.1560235

84. Hung A, Kim YH, Pavon JM. Deprescribing in older adults with polypharmacy. BMJ. 2024 May 7;e074892. doi:10.1136/bmj-2023-074892

85. Hawiger J, Niewiarowski S, Gurewich V, Thomas DP. Measurement of fibrinogen and fibrin degradation products in serum by staphylococcal clumping test. J Lab Clin Med. 1970 Jan;75(1):93–108. PubMed PMID: 4243578.

86. Wiegand I, Hilpert K, Hancock REW. Agar and broth dilution methods to determine the minimal inhibitory concentration (MIC) of antimicrobial substances. Nat Protoc. 2008 Feb 17;3(2):163–75. doi:10.1038/nprot.2007.521

87. Odds FC. Synergy, antagonism, and what the chequerboard puts between them. Journal of Antimicrobial Chemotherapy. 2003 Jun 12;52(1):1–1. doi:10.1093/jac/dkg301

88. Buttress JA, Halte M, te Winkel JD, Erhardt M, Popp PF, Strahl H. A guide for membrane potential measurements in Gram-negative bacteria using voltage-sensitive dyes. Microbiology (N Y). 2022 Sep 30;168(9). doi:10.1099/mic.0.001227

89. Borrelli C, Douglas EJA, Riley SMA, Lemonidi AE, Larrouy-Maumus G, Lu WJ, et al. Polymyxin B lethality requires energy-dependent outer membrane disruption. Nat Microbiol. 2025 Sep 29;10(11):2919–33. doi:10.1038/s41564-025-02133-1

