## Supplementary Figures for "Commonly prescribed medicines antagonise anti-MRSA antibiotics and select for resistance"

### Supplementary Data file

**Supplementary Table. 1. Commonly used medications in the UK.** Medications are categorised by class, indication, and number of annual UK prescriptions. Prescription data was obtained from the NHS Healthcare and & Prescribing Data, available at prescribemap.com.

| Medication | Class | Indication | Number of prescriptions UK (in millions) |
|---|---|---|---|
| Simvastatin | Statin | Hypercholesterolemia; prevention of cardiovascular disease | 11.7 |
| Fluoxetine | Selective serotonin reuptake inhibitor | Major depressive disorder, anxiety disorder | 7.5 |
| Amlodipine | Calcium channel blocker | Hypertension and angina | 39.8 |
| Levothyroxine | Thyroid hormone replacement | Hypothyroidism | 34.5 |
| Furosemide | Loop diaretic | Oedema and hypertension | 11.2 |
| Prednisolone | Corticosteroid | Inflammatory and autoimmune conditions | 6.4 |
| Metformin | Biguanide | Type 2 diabetes mellitus | 26.6 |
| Omeperazole | Proton pump inhibitor | Gastroesophageal reflux disease, peptic ulcers | 35.8 |

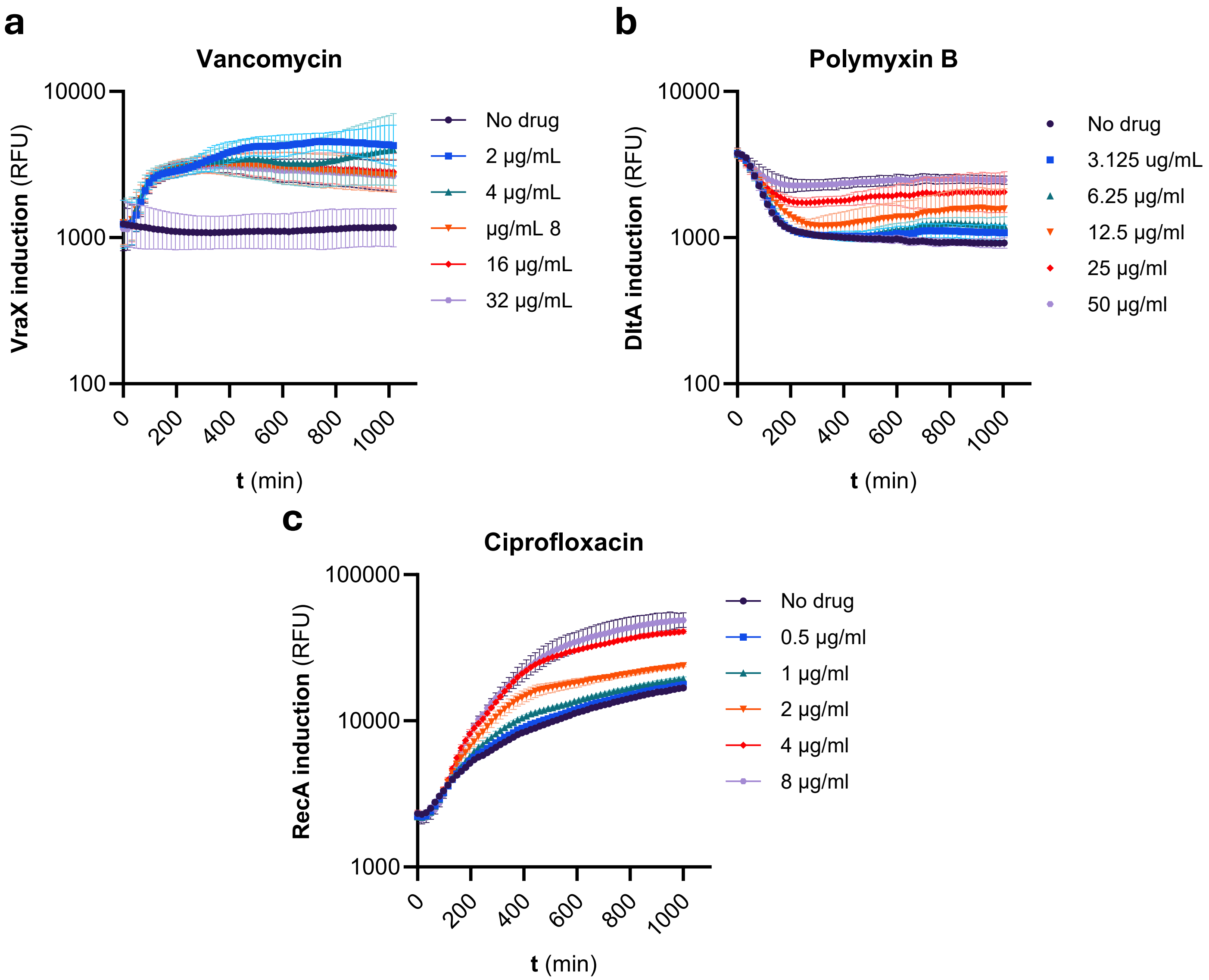


**Supplementary Figure 1. GFP-fusion plasmids are induced by appropriate antibiotic controls. (a)** Induction assay of p*vraX*-GFP in the presence of a 1:2 dilution series of vancomycin as determined by GFP accumulation over time. **(b)** Induction assay of p*dltA*-GFP in the presence of a 1:2 dilution series of polymyxin B as determined by GFP accumulation over time. **(c)** Induction assay of p*recA*-GFP in the presence of a 1:2 dilution series of ciprofloxacin as determined by GFP accumulation over time. All experiments were replicated in n=3 independent assays. Error bars show the standard deviation of the mean**.**

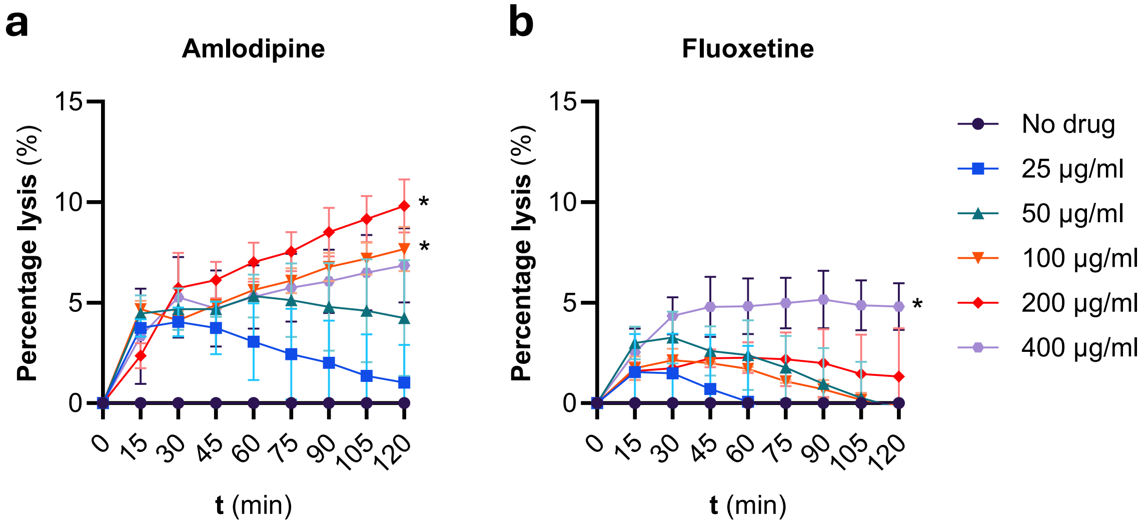


**Supplementary Figure 2. Amlodipine and fluoxetine induce bacterial lysis. (a)** Percentage lysis of *S. aureus* JE2 over time, exposed to a 1:2 dilution series of amlodipine. **(b)** Percentage lysis of *S. aureus* JE2 over time, exposed to a 1:2 dilution series of fluoxetine. All experiments were replicated in n=3 independent assays. Error bars show the standard deviation of the mean**.** Significance differences were determined between the no drug condition and the 400 µg/ml and 200 µg/ml treated conditions by two-way repeated measures ANOVA with post hoc Dunnett's test to correct for multiple comparisons. *$P < 0.05$.

**Supplementary Table. 2. NTML strains associated with reduced susceptibility to simvastatin.**

Strains are categorised by gene name (where available), indication, and simvastatin MIC.

| Strain reference | Gene name | Function | MIC (µg/mL) |
|---|---|---|---|
| NE37 | *icaA* | Biofilm | 50 |
| NE234 | *cap5I* | Capsule | 50 |
| NE75 | *cap1B* | Capsule | 100 |
| NE1495 | *murA* | Cell wall | 50 |
| NE1693 | *yycH* | Cell wall | 50 |
| NE267 | *sgtA* | Cell wall | 100 |
| NE596 | *sgtB* | Cell wall | 200 |
| NE217 | *pknB* | Cell wall | 200 |
| NE1369 | *lytH* | Cell wall | 200 |
| NE1713 | *alr* | Cell wall | 100 |
| NE1099 | *lyrA* | Cell wall | 50 |
| NE945 | *brnQ* | Central metabolism | 50 |
| NE955 | *narI* | Central metabolism | 50 |
| NE1717 | *aroC* | Central metabolism | 50 |
| NE476 | *fba* | Central metabolism | 50 |
| NE198 | *ald* | Central metabolism | 50 |
| NE232 | | Central metabolism | 50 |
| NE5 | | Central metabolism | 50 |
| NE6 | | Central metabolism | 50 |
| NE233 | *glpD* | Central metabolism | 50 |
| NE1896 | *ipdA* | Central metabolism | 100 |
| NE16 | *moaD* | Central metabolism | 200 |
| NE477 | *deoD* | DNA replication/ repair/ nucleotide metabolism | 50 |
| NE1390 | | DNA replication/ repair/ nucleotide metabolism | 50 |
| NE243 | *polA* | DNA replication/ repair/ nucleotide metabolism | 50 |
| NE246 | | DNA replication/ repair/ nucleotide metabolism | 50 |
| NE242 | *dprA* | DNA replication/ repair/ nucleotide metabolism | 50 |
| NE277 | *tdk* | DNA replication/ repair/ nucleotide metabolism | 100 |
| NE1541 | | DNA replication/ repair/ nucleotide metabolism | 100 |
| NE279 | | DNA replication/ repair/ nucleotide metabolism | 100 |
| NE1744 | | Hypothetical | 50 |
| NE18 | | Hypothetical | 50 |
| NE1402 | | Hypothetical | 50 |
| NE721 | | Hypothetical | 50 |
| NE1520 | | Hypothetical | 50 |
| NE1372 | | Hypothetical | 50 |
| NE257 | | Hypothetical | 50 |
| NE230 | | Hypothetical | 50 |
| NE4 | | Hypothetical | 50 |
| NE38 | | Hypothetical | 50 |
| NE40 | | Hypothetical | 50 |
| NE1809 | | Hypothetical | 100 |
| NE1861 | | Hypothetical | 50 |
| NE1892 | | Hypothetical | 50 |
| NE1831 | | Hypothetical | 50 |
| NE268 | | Hypothetical | 100 |
| NE53 | | Hypothetical | 50 |
| NE1800 | | Hypothetical | 100 |
| NE1585 | | Hypothetical | 100 |
| NE265 | | Hypothetical | 100 |
| NE1795 | | Hypothetical | 100 |
| NE1703 | | Hypothetical | 100 |
| NE50 | | Hypothetical | 50 |
| NE1734 | | Hypothetical | 200 |
| NE1909 | | Hypothetical | 200 |
| NE208 | | Hypothetical | 200 |
| NE86 | | Hypothetical | 100 |
| NE87 | | Hypothetical | 100 |
| NE258 | *cls* | Membrane | 100 |
| NE209 | | Phage | 50 |
| NE244 | | Phage | 50 |
| NE699 | *clpC* | Protein synthesis | 50 |
| NE289 | | Protein synthesis | 50 |
| NE195 | *pepF* | Protein synthesis | 50 |
| NE2 | | Protein synthesis | 50 |
| NE896 | *spxH* | Protein synthesis | 50 |
| NE1752 | *rbfA* | Protein synthesis | 50 |
| NE1662 | *miaB* | Protein synthesis | 200 |
| NE1269 | *ohr* | Redox | 50 |
| NE1345 | *menD* | Redox | 50 |
| NE1669 | *nreC* | Redox | 50 |
| NE1596 | *bshB2* | Redox | 50 |
| NE281 | | Transcription regulator | 100 |
| NE197 | | Transport | 50 |
| NE269 | | Transport | 100 |
| NE13 | | Transport | 200 |
| NE1890 | | Virulence | 100 |
